# A protocol for the use of cold plasma treatment to inhibit *in vitro* growth of *Fusarium graminearum*

**DOI:** 10.1101/2022.05.23.493010

**Authors:** M. Kaur, D. Hüberli, K. L. Bayliss

## Abstract

Cold plasma is an ionised gas that can be used to control a range of microorganisms. In this study, a protocol was developed for cold plasma treatment of *in vitro* growth of *Fusarium graminearum*, a global mycotoxic species generally associated with cereal grain. Four *F. graminearum* isolates on potato dextrose agar (PDA) were treated with cold plasma for 70 s from a distance of 21 cm in a closed environment, and their radial colony growth was measured. To consider whether cold plasma modified the culture media, non-inoculated half-strength PDA plates were also treated with cold plasma prior to inoculation with the *F. graminearum* isolates. Similarly, to determine if a rise in temperature during the treatment impacted the growth of the isolates or the culture media itself, the plates were treated with dry heat before and after inoculation with the isolates. Treatment of *F. graminearum* isolates with cold plasma inhibited their growth and was not associated with the culture media or the rise in temperature during the treatment. Optical emission spectroscopy of cold plasma identified reactive (ionised) species of argon, nitrogen, hydrogen, oxygen, copper, and carbon with the highest number of peaks produced for argon. These results demonstrate that cold plasma can significantly reduce the *in vitro* growth of *F. graminearum* isolates when treated in a closed environment and suggest there is potential to control the *in vivo* growth of *F. graminearum*.

## Introduction

*Fusarium graminearum* is a devastating fungal pathogen associated with fusarium head blight (FHB) in wheat and other cereal crops. *F. graminearum* chlamydospores and perithecia survive in host crop residues or stubble and serve as the primary source of inoculum (Wegulo, 2012). Ascospores or conidia are released from the infected stubble under warm wet conditions, and dispersed to nearby wheat plants by wind or rain splash, where they germinate and infect susceptible wheat heads, initiating the development of FHB (Wegulo, 2012). An integrated management approach to reduce inoculum levels through crop rotation and fungicide application during or before anthesis (approx. GS 59 - 65) is the primary strategy to mitigate disease development (Blandino et al., 2012; Wegulo et al., 2011). However, the integrated management strategy’s efficacy depends on many factors such as cultivar resistance to the pathogen, weather conditions, and the type of fungicide used (Blandino et al., 2012; Edwards, 2004). Furthermore, if the disease infection occurs at later stages of grain development, grain may appear normal and potentially carry the pathogen to storage (Osborne & Stein, 2007) intensifying the need for an effective and reliable treatment.

Mycotoxins, such as trichothecenes, are also produced by *F. graminearum*. These are toxic secondary metabolites, and the consumption of products from mycotoxin-contaminated cereal grain causes serious health issues in humans and animals (Foroud & Eudes, 2009). Trichothecenes are mainly produced during crop maturation or when the grain are stored under high humidity and temperature (Hope et al., 2005). Once the mycotoxins are established in cereal grain, they are difficult to eliminate due to their chemical and thermal stability (Generotti et al., 2015). Cold plasma could potentially provide a solution for managing postharvest problems of *F. graminearum* in cereal grain.

Cold plasma is an ionised form of gas, which contains reactive oxygen species (ROS), reactive nitrogen species (RNS), electrons, and other free radicals. ROS and RNS are known to damage microbial cells through the oxidation of cytoplasmic membranes, lipids, proteins, and DNA strands (Bourke et al., 2017; Misra et al., 2019), which induce membrane perforation and other changes in the hyphal surface, thereby impacting growth (Avramidis et al., 2010; Simoncicova et al., 2018). The potential of cold plasma to reduce seed-borne fungal contamination in wheat, oats, corn, and other grain was first reported by Selcuk et al. (2008). They demonstrated that up to a 99 % reduction in fungal load of *Aspergillus* and *Penicillium* species on wheat grain was achievable following cold plasma treatment for 15 min; however, disinfection was dependent on the gas used to generate the plasma and the surface of the treated grain, amongst other factors. Other authors have reported similar findings with different grain commodities, including rice (Kang et al., 2015; Ochi et al., 2017) and barley (Los et al., 2018).

A problem with most of the research in this field is that published studies have used widely different and often custom-built plasma generation systems, resulting in a diverse range of treatments that make comparisons difficult and identification of optimal conditions almost impossible (Sarangapani et al., 2018). The effectiveness of cold plasma can be influenced by factors such as the electrical voltage used to generate the plasma and the duration of treatment of the commodity with the plasma (Braşoveanu et al., 2015). Therefore, it is essential to define each commodity’s treatment conditions or microbe of interest based on the cold plasma equipment being used.

This study developed optimised methods for cold plasma treatment of *in vitro* cultures of *F. graminearum.* Experiments were also designed to test the hypothesis that cold plasma inhibits the *in vitro* growth of *F. graminearum* cultures isolated from postharvest wheat and barley.

## Materials and Methods

### Fungal cultures

Four *F. graminearum* isolates, WAC11354, WAC11387, WAC11490 and WAC12336, were provided by the Department of Primary Industries and Regional Development (DPIRD) in Western Australia. Of these isolates, two were isolated from wheat grain, one from barley grain and one from wheat stubble (Table 1). Two isolates, WAC11354 and WAC11490, were associated with FHB disease in wheat (previously determined by DPIRD). The isolates were grown on half-strength Potato Dextrose Agar (PDA [Difco] powder 19.5 g + Agar [Difco] 9.5 g + deionised water 1 L, referred to as ½ PDA for all experiments), in the dark at 25±1°C. Disks of 3 mm taken from the edge of actively growing, four-day-old cultures were aseptically transferred to 10 mL of ½ PDA in 90 mm Petri plates immediately before cold plasma treatment.

**Table 1.** *Fusarium graminearum* isolates used for all the treatments, their host species, collection year and the place of origin.

| WAC<sup>ab</sup> number | Host species | Collection year | Isolated from |
| --- | --- | --- | --- |
| WAC11354 | *Triticum aestivum* (Wheat) | 2005 | Grain |
| WAC11387 | *T. aestivum* | 2004 | Grain |
| WAC11490 | *T. aestivum* | 2004 | Stubble |
| WAC12336 | *Hordeum vulgare* (Barley) | 2004 | Grain |

<sup>a</sup> WAC-WA Culture Collection numbers allocated by the Department of Primary Industries and Regional Development Herbarium.
<sup>b</sup> Locality of origin of all the isolates- Wellstead, Western Australia.

### Cold plasma Equipment

A Blown-Arc Pro Series plasma surface treater (Enercon, Wisconsin, US) was used for all experiments. The Blown-Arc Pro produces a gliding arc plasma with a maximum output voltage of 3.4 KV_RMS_ at an output frequency range of 20-60 kHz. The plasma is generated by supplying compressed atmospheric air at 70-90 psi through high voltage electrodes.

### Optical emission spectroscopy and temperature measurement

A fibre optic spectrometer (Avantes AvaSpec-2048-8 with AvaSpec v. 8.3.1.0 software) was used to monitor the reactive oxygen and nitrogen species produced during plasma treatment. A 400 nm fibre optic cable was placed at a distance of 21 cm from the plasma emission point to detect the spectra of reactive species, as previously described by Siddique et al. (2019). The Blown-Arc Pro emission spectra were measured at the centre and edge of the plasma flame and then combined to produce a final spectrum for each environment.

The temperature during treatment was recorded every 15 s by placing a four-inch food-grade stainless steel probe attached to a Hobo UX120-006 4-Channel Analog Logger (Temperature Sensor TMC6-HC) at the level of the Petri plate surface.

### Optimised cold plasma treatment of F. graminearum

Following a series of preliminary trials (data not shown), the Blown-Arc Pro was optimised to treat the *F. graminearum* isolates for a duration of 70 s from a distance of 21 cm inside a closed box which was referred as a closed environment treatment, with the Petri plate lids removed. All plates were sealed with plastic food wrap (Glad^®^ Wrap) and incubated at 25±1°C in the dark after treatment. The radial colony growth was measured every 24 hr by recording the cultures’ diameter on the two perpendicular axes for up to five days or until the untreated control cultures reached the edge of the plate. The radial colony growth was then calculated as the average growth rate over the five days. There were ten replicate plates for each isolate, and the entire trial was conducted twice.

### Effect of media treated with cold plasma on F. graminearum growth

To determine whether cold plasma treatment modified the culture media and consequently inhibited culture growth, non-inoculated ½ PDA was also treated with cold plasma with the Petri plate lids removed, using the optimised conditions described above. Treated PDA was then aseptically inoculated with disks of actively growing cultures of all four isolates and radial colony growth was measured as described above. There were ten replicates, and this trial was conducted twice.

### Effect of heat generated during cold plasma treatment on F. graminearum growth

While conducting the preliminary trials, the cold plasma produced by the Blown-Arc Pro was observed to increase the temperature at the surface of the PDA plate to no more than 120 °C in the closed environment treatment (data not shown). Therefore to eliminate any possible impact of temperature on culture growth or directly on PDA during treatment, two dry heat trials were conducted. In the first trial, disks of 3 mm taken from the edge of actively growing, four-day-old cultures were aseptically transferred to 10 mL of ½ PDA in 90 mm Petri plates as described above, sealed with plastic food wrap, and placed in an oven at 120±2 °C for 80 s. In the second trial, non-inoculated ½ PDA plates were placed in the oven as described above and then aseptically inoculated. Controls in both trials were kept at room temperature. All cultures were incubated at 25±1 °C in the dark, and radial colony growth was measured as described above. There were ten replicate plates for each isolate in each trial, and both trials were conducted twice.

### Statistical analysis

All data were expressed as means and standard errors of all replicates. The repeated measure ANOVA showed no significant difference between the repeated trials; therefore, all means of the repeated trials were combined. The data were analysed using One-way Analysis of Variance (ANOVA) and compared by Fisher’s Least Significant Difference (LSD) in RStudio (version 3.5.2). The level of significance was considered below 0.05. The results were plotted using ggplot packages in R.

## Results

### Impact of cold plasma on F. graminearum growth rates

The application of cold plasma to *in vitro* cultures of *F. graminearum* for 70 s from a distance of 21 cm significantly (P < 0.001) reduced the growth rate of all isolates in a closed environment, although WAC11387 was less affected than the other isolates (Fig. 1a). Further, the treatment of ½ PDA with cold plasma for 70 s before inoculation showed no noticeable (P > 0.2) impact on the media, such that it subsequently influenced *F. graminearum* growth rates. However, the treatment had a significant interaction (P = 0.01, Fig. 1b) with the isolates. At the individual isolate level, the isolate WAC12336 had a significantly lower growth rate on treated ½ PDA plates compared to untreated plates, whereas the growth rate of the other three isolates did not change significantly from the controls.

**Fig. 1.**
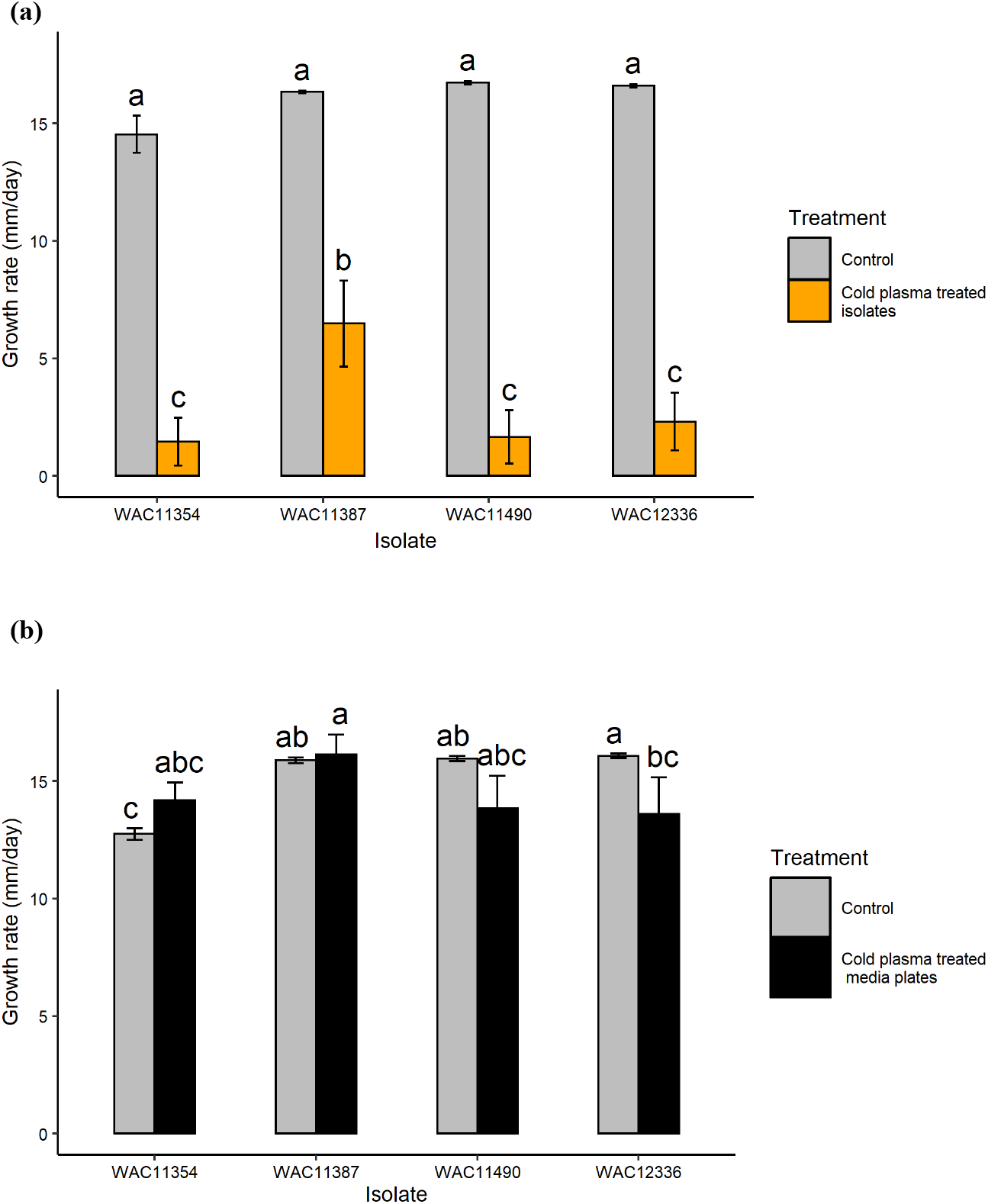
Cold plasma treatments of four *Fusarium graminearum* isolates for 70 s at a distance of 21 cm from the plasma emission point in a closed environment with the Blown-Arc Pro. The graphs represent a) cold plasma treatment of inoculated plates and b) cold plasma treatment of half-strength Potato Dextrose Agar plates that were then immediately inoculated. Bars = standard errors of the mean (n = 20). Means with the same letter are not significantly different according to Fisher’s Least Significant Difference test.

Dry heat treatment of plates inoculated with *F. graminearum* isolates (Fig. 2a) also had no significant (P = 0.497) influence on culture growth rate. Similarly, when the PDA was treated with dry heat and inoculated immediately after the treatment, no significant (P = 0.167) impact of the treatment was observed for the isolates grown on the treated plates (Fig. 2b). Therefore, the rise in temperature during cold plasma treatment did not influence the growth rate of *F. graminearum*.

**Fig. 2.**
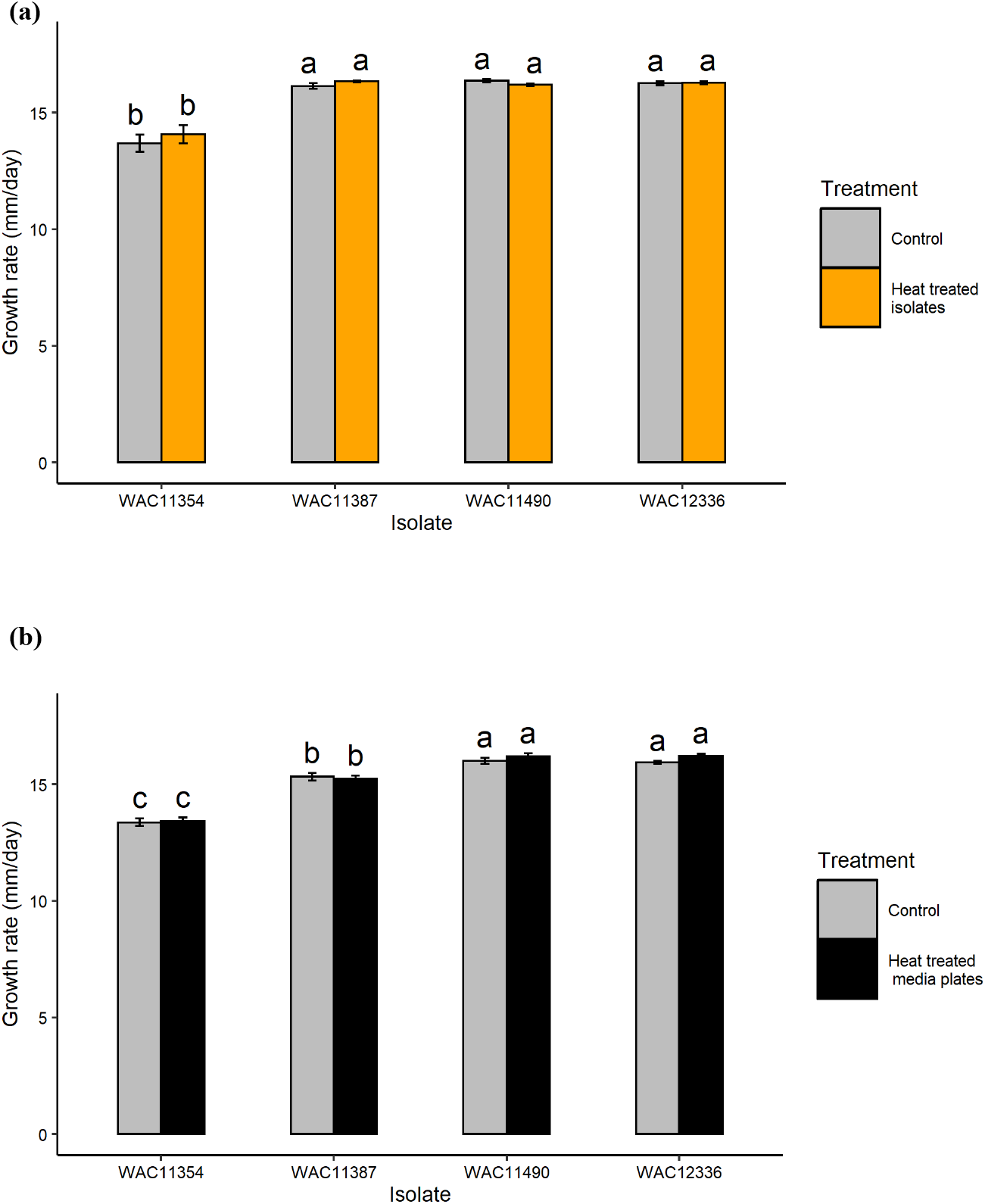
Dry heat treatments of four *Fusarium graminearum* isolates performed at 120 °C for 80 s. The graphs represent a) dry heat treatment of inoculated plates and b) dry heat treatment of half-strength Potato Dextrose Agar plates that were immediately inoculated. Bars = standard errors of the mean (n = 20). Means with the same letter are not significantly different according to Fisher’s Least Significant Difference test.

### Optical emission spectroscopy of plasma generated by the Blown-Arc Pro

The Blown-Arc Pro produced a total of 28 peaks for different ionised molecules. At a selected noise/signal level of 15, the peak number and the intensity of the peaks produced by the Blown-Arc Pro were highest for argon (Ar) (Fig. 3). High numbers and intensities of peaks were also observed for ionised nitrogen (N), oxygen (O), copper (Cu), and carbon (C II). Other molecules detected by the emission spectra were ionised carbon dioxide (CO_2_^+^), carbon monoxide anion (CO^+^), dihydrogen (H_2_), and amino radical (NH_2_), dinitrogen (N_2_), N_2_^+^, hydroxide (OH), and oxygen molecules (O_2_), cyclopropatriene (C_3_) and nitrogen dioxide (NO_2_).

**Fig. 3.**
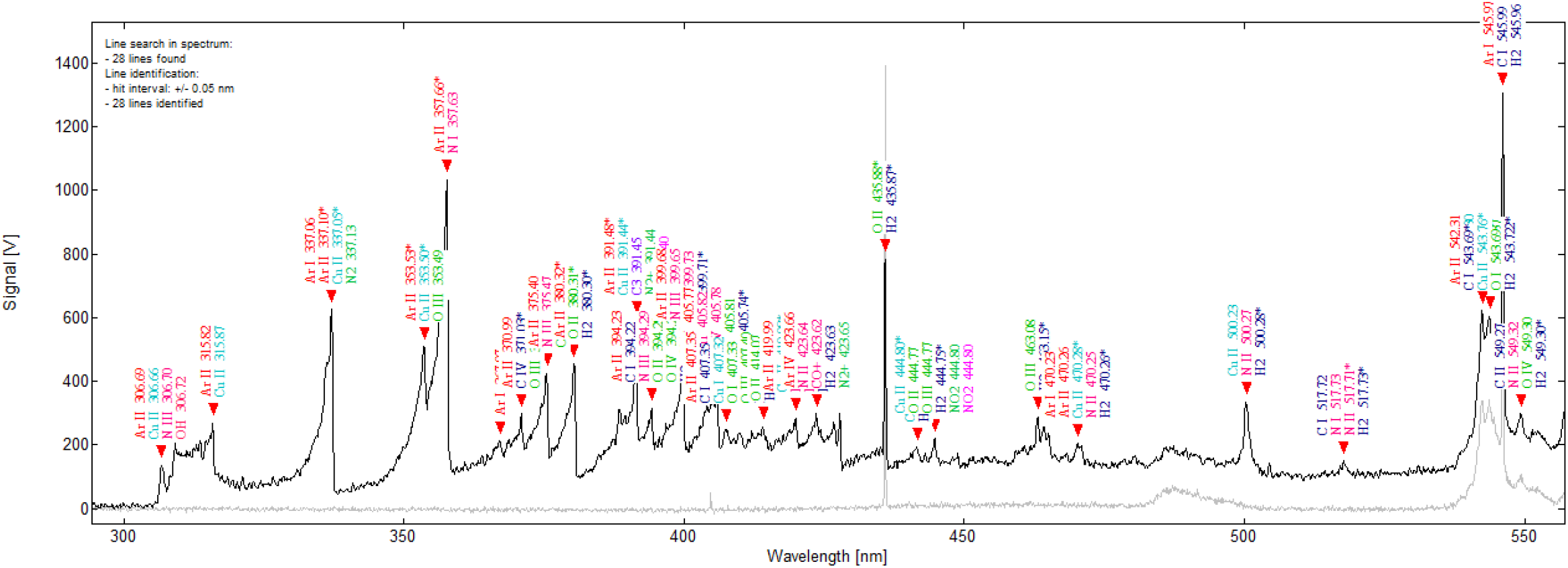
The spectra of reactive species produced by Blown-Arc Pro in a closed environment. The grey line on the graphs represents the reference spectra produced from background light when the Blown-Arc Pro was off, whereas the black line shows the spectra of actively producing cold plasma when the Blown-Arc Pro was running. The ionisation states of reactive oxygen (O), Nitrogen (N), Carbon (C), Argon (Ar) and Copper (Cu) given in spectroscopic notation denotes: ‘I’-neutrals, ‘II’-single ionised elements, ‘III’-double ionised elements, and ‘IV’-triple ionised elements.

## Discussion

This study has demonstrated that cold plasma treatment can significantly reduce the *in vitro* growth rate of *F. graminearum* colonies. This finding was consistent with Na et al. (2013), who demonstrated that *in vitro* cold plasma treatment of 60 s or longer completely inhibited the spore germination and hyphal growth of *F. graminearum*. In contrast to Na et al. (2013), the current study also demonstrated that the temperature generated during treatment and the impact of cold plasma on the culture medium did not influence culture growth. Conclusively, the inhibition of *F. graminearum* growth was a result of cold plasma treatment.

The closed treatment environment chosen for the current study seems to be commonly practised for cold plasma application to different microorganisms and commodities. The use of cold plasma in a closed or sealed environment has been demonstrated to successfully treat various bacterial and fungal pathogens (Los et al., 2018; Los et al., 2020; Misra et al., 2014; Ouf et al., 2015; Selcuk et al., 2008) and food commodities (Ouf et al., 2015; Selcuk et al., 2008). It was suggested by Siddique et al. (2019) that there is higher efficacy of the treatment in a closed environment due to the higher intensity of ionised molecules observed when compared to an open environment. The ionised molecules are likely dissipated into the air during an open environment treatment. Interestingly, Siddique et al. (2019), using a similar type of gliding arc plasma generator (Enercon Dyne-A-Mite), only observed three peaks during cold plasma treatment in the closed environment. In contrast, in the current study, the Blown-Arc Pro produced 28 peaks, indicating distinct differences between similar forms of cold plasma. The properties of the cold plasma produced by different machines should be considered when developing a pathogen treatment protocol.

Currently, the grain industry prominently relies on foliar fungicides to manage the pathogen in the field (Wegulo et al., 2015) in addition to other management practices. However, the efficacy of these fungicides highly depends upon the timing of the fungicide application, coverage, crop cultivars and virulence of isolates (Mesterházy et al., 2003). Becher et al. (2010) reported that the *F. graminearum* strains can adapt to these fungicides, potentially developing completely new isolates with stronger virulence. The emission spectroscopy in the current study showed that cold plasma produced by the Blown-Arc Pro contains various ionised molecules of N, O and C which could potentially be responsible for the inhibition of *F. graminearum* growth. Though the mode of action of cold plasma yet requires further investigation, other authors have found that plasma reactive species changes the hyphae surface and spores’ structure thus reducing the fungi’s ability to grow (Kang et al., 2015; Simoncicova et al., 2018). It also kills pathogens by damaging microbial cell structures such as cell membranes, lipids, proteins, and DNA strand via ROS and RNS (Bourke et al., 2017). Due to its multimodal and non-specific action mechanism, it is unlikely that the pathogen will adapt or develop resistance against the treatment. It is also evident from the study that the efficacy of the treatment is consistent against multiple isolates of *F. graminearum*. Therefore, cold plasma could provide an advantageous solution for the global grains industry.

A possible application of cold plasma for the grains industry could be treating seed grain to prevent seedling blight caused by *F. graminearum*. Seedling blight primarily originates from planting *Fusarium* infected seed which carries the pathogen but are asymptomatic (Jones, 1999). Gilbert et al. (2003) demonstrated that the infected seed impacts seed germination up to 75 % possibly by overtaking and killing the seed at germination, thus ultimately reducing crop productivity. Under warm and humid conditions, the fungus from infected seed could produce prithecia and wind-spread ascospores, and increase the FHB severity (Gilbert et al., 2003). Apart from fungicide seed treatment, currently no other treatments are available to manage *F. graminearum* infected seed. Similar to foliar fungicides, as stated above, the efficacy of fungicide seed treatment depends on the inoculum level present on the seed and the environmental conditions to develop seedling blight (Hysing & Wiik, 2014). A potential use of cold plasma treatment could be to reduce the fungal load present on the infected seed, which would be investigated in future.

In conclusion, this study confirmed that cold plasma could significantly reduce the *in vitro* growth of *F. graminearum* in a closed environment. Cold plasma does not influence the pathogen growth due to its impact on culture media or rise in the temperature during the treatments. The emission spectroscopy showed the presence of different ionised molecules in the cold plasma generated during the treatment. However, the study also indicated that the intensity of these ionised molecules could differ with the type of machine used to generate the plasma. For this reason, it is essential to standardise the cold plasma treatment conditions, to effectively control plant pathogens before beginning trials on an industrial scale.

## Acknowledgments

The first author gratefully acknowledges the financial support provided by the Department of Primary Industries and Regional Development under the Boosting Grains, Research and Development Postgraduate Scholarships program. The author is also thankful for the feedback and valuable advice from Prof. Jen McComb and Prof. Giles Hardy, who helped improve this study’s context.

## Author information

### Affiliations

**College of Science, Health, Engineering and Education, Murdoch University, 90 South Street, Murdoch WA 6150, Australia**

Maninder Kaur

**College of Science, Health, Engineering and Education, Murdoch University, 90 South**

**Street, Murdoch WA 6150, Australia**

Kirsty L. Bayliss

**Food Futures Institute, Murdoch University, 90 South Street, Murdoch WA 6150, Australia**

Kirsty L. Bayliss

**Department of Primary Industries and Regional Development, 3 Baron-Hay Court, South Perth WA 6151, Australia**

Daniel Hüberli

### Corresponding author

Correspondence to Kirsty L. Bayliss

### Author contributions

All authors contributed to the study’ s conception and design. Material preparation, data collection and analysis were performed by Maninder Kaur. The first draft of the manuscript was written by Maninder Kaur and all authors commented on previous versions of the manuscript. All authors read and approved the final manuscript.

## Declarations

### Availability of data and material (data transparency)

The current study data are not publicly available due to the technology’s commercial interest but are available from the corresponding author on reasonable request.

### Code availability

All data analysis was performed using Rstudio (version 3.5.2) and the codes available upon request.

### Ethical approval

This article does not contain any studies with human participants or animals performed by any of the authors.

